# Epigenetic Programming of Macrophage Phenotypes by STING–IRF3 Drives Inflammation in Ascending Thoracic Aortic Dissection

**DOI:** 10.64898/2026.01.22.701198

**Authors:** Bowen Li, Chen Zhang, Samantha Xu, Yanming Li, Deborah Vela, Hernan Vasquez, Lin Zhang, Abhijit Chakraborty, Hong S Lu, Joseph S Coselli, Toru Suzuki, Alan Daugherty, Dianna M. Milewicz, Ziad Mallat, Liwu Li, Scott A LeMaire, Ying H Shen

**Author notes:** These authors have equal contributions to this manuscript (Bowen Li, Chen Zhang, Samantha Xu, Yanming Li). Correspondence to: Ying H. Shen, MD, PhD, Michael E. DeBakey Department of Surgery, Baylor College of Medicine,; Scott A. LeMaire, MD, Geisinger, 100 N. Academy Avenue, Danville, PA 17822-4400.

## Abstract

**Background:** Ascending thoracic aortic dissection (ATAD) is characterized by extensive macrophage (MΦ) accumulation and profound inflammation; however, the mechanisms sustaining pro-inflammatory MΦ activation remain incompletely defined. Emerging evidence indicates that epigenetically generated immune memory drives innate immune cells toward persistent inflammatory states. In this study, we investigated whether epigenetic reprogramming governs MΦ phenotypic fate and contributes to ATAD pathogenesis.

**Methods:** We performed single-cell RNA sequencing of human ascending aortic tissues from controls, patients with ascending thoracic aortic aneurysm (ATAA), and patients with acute ascending thoracic aortic dissection (ATAD). We also performed integrated single-cell RNA sequencing, single-cell ATAC sequencing, and spatial transcriptomics in an angiotensin II (Ang II)–infused mouse model. The role of the STING–IRF3 signaling axis in MΦ epigenetic programming was examined using MΦ-*Sting ^-/-^* and MΦ-*Irf3^-/-^* mice.

**Results:** In human and mouse aortic tissues, we identified multiple functional MΦ populations including pro-inflammatory, phagocytic/anti-inflammatory, proliferative, and reparative/healing MΦs. Aortic MΦs in both sporadic ATAD patients and Ang II–induced ATAD mice exhibited a pronounced pro-inflammatory bias with enhanced differentiation toward pro-inflammatory MΦs and impaired differentiation toward phagocytic/anti-inflammatory states. Pro-inflammatory MΦs were particularly abundant in dissection sites, whereas phagocytic MΦs were enriched in discrete adventitial niches. Origin analyses revealed a substantial increase in *CCR2*⁺ recruited MΦs within the aortic wall, which preferentially differentiated into pro-inflammatory MΦs. In contrast, *LYVE1*⁺ resident MΦs— predominantly biased toward phagocytic phenotypes—were markedly depleted in ATAD. Single-cell ATAC sequencing identified coordinated chromatin remodeling with increased accessibility at pro-inflammatory gene loci and decreased accessibility at phagocytic gene loci. Among candidate transcriptional regulators identified, IRF family TFs, including IRF3 emerged as unique factors capable of simultaneously promoting pro-inflammatory gene programs while suppressing phagocytic gene expression.

Mechanistically, STING–IRF3 signaling orchestrates this biased transcriptional state, likely through coordinated BRG1-dependent chromatin opening at pro-inflammatory gene loci and chromatin closing at phagocytic/anti-inflammatory gene loci. MΦ specific *Sting ^-/-^* and *Irf3^-/-^* mice exhibited attenuated inflammatory reprogramming and reduced aortic destruction and dissection.

**Conclusions:** These findings identify STING–IRF3–mediated epigenetic programming of MΦs as a fundamental mechanism driving aortic inflammation and ATAD development. Targeting MΦ epigenetic programming may represent a promising therapeutic strategy to prevent aortic dissection.

**Graphic Abstract:** 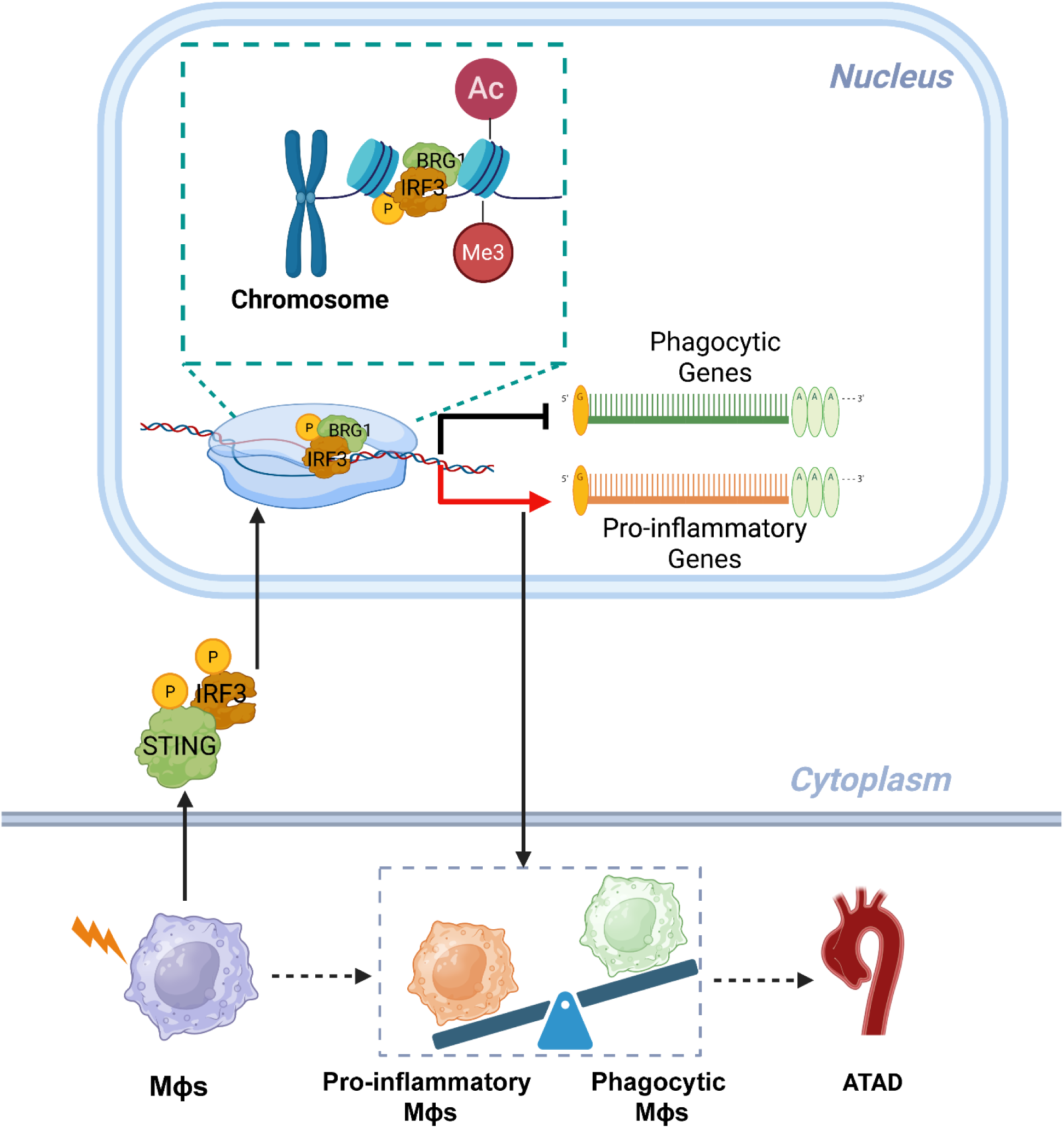

## INTRODUCTION

Sporadic ascending thoracic aortic aneurysm (ATAA) is a cardiovascular condition characterized by the progressive loss of integrity of the aortic wall^1–3^. Without surgical treatment, ATAA often progresses to acute ascending thoracic aortic dissection (ATAD) and its lethal consequences, including rupture and pericardial tamponade. Unfortunately, there are currently no pharmacological treatments that reliably prevent disease progression from ATAA to ATAD^4,5^. Therefore, there is an urgent need to understand the mechanisms underlying ATAD development to develop effective therapeutic strategies to prevent ongoing aortic degeneration and eventual catastrophic tissue failure.

One of the defining features of ATAD is profound and persistent inflammation, characterized by marked accumulation of inflammatory cells—particularly macrophages (MΦs)^6,7^. Despite its central role in disease progression, the mechanisms underlying this unresolved inflammatory state remain poorly understood. MΦs are highly plastic immune cells with diverse functions. Beyond promoting inflammation and tissue destruction, MΦs also clear cellular debris, orchestrate resolution of inflammation, and support tissue repair. These diverse functions are mediated by distinct MΦ subpopulations with specialized phenotypes^8–11^. However, how these heterogeneous MΦ populations dynamically evolve during ATAD development remains unclear.

From a developmental and ontogenetic perspective, MΦs can be broadly categorized as tissue-resident cells or recruited cells. Resident MΦs arise from yolk sac and fetal liver progenitors during embryogenesis and persist within specialized tissue niches, whereas recruited MΦs originate as bone marrow–derived monocytes that infiltrate tissues in response to injury or inflammation^12,13^. The dynamics of resident versus recruited MΦs during ATAD progression, and whether each population undergoes distinct differentiation trajectories into functional subtypes within the aortic wall, remain largely undefined.

Although prior studies in aortic disease have defined pro- and anti-inflammatory MΦ phenotypes and highlighted roles for cytokines and chemokines^7^, the upstream mechanisms governing MΦ phenotypic acquisition, maintenance, and diversification in the diseased aorta remain poorly understood. Emerging evidence shows that innate immune cells, including monocytes and MΦs, can undergo trained immunity, a form of epigenetically mediated immune memory that induces long-lasting functional reprogramming in response to prior stimuli^14–17^. This training can occur in the bone marrow or peripheral tissues and is shaped by the nature, strength, and duration of the initiating signals. Dysregulated trained immunity has been implicated in chronic inflammatory cardiovascular diseases characterized by persistent and exaggerated inflammation^15,18^. Whether trained immunity contributes to the differentiation of MΦs toward pro-inflammatory phenotypes in ATAD—thereby sustaining chronic aortic inflammation—remains unknown. Moreover, the identity of the initiating stimuli, as well as the key transcription factors and epigenetic regulators that drive maladaptive pro-inflammatory training in the aortic wall, have yet to be defined.

In the present study, we combined single-cell multi-omics, spatial transcriptomics, and functional in vitro and in vivo approaches to define the phenotypic landscape, origin, and regulatory mechanisms of aortic MΦs during ATAD development. We identified a shift from phagocytic toward pro-inflammatory MΦ phenotypes in both human ATAD tissues and an angiotensin II (Ang II)–induced mouse ATAD model. Integrated single-cell chromatin accessibility profiling revealed that this phenotypic modulation is driven by epigenetic remodeling, characterized by increased chromatin accessibility at inflammatory gene loci and repressive chromatin remodeling at phagocytic gene loci. Mechanistically, we uncovered the STING–IRF3 signaling as a critical upstream pathway orchestrating this epigenetic reprogramming. This pathway recruits the chromatin remodeler BRG1 to pro-inflammatory gene loci, inducing permissive chromatin states. Genetic deletion of *Sting* or *Irf3* specifically in MΦs attenuated MΦ inflammatory reprogramming and reduced ATAD formation in vivo.

Together, these findings identify epigenetic reprogramming of MΦ phenotypes through STING–IRF3 signaling as a fundamental mechanism driving aortic inflammation and ATAD progression and point to this pathway as a promising therapeutic target.

## METHODS

### Data Availability

All detailed methods are presented in the Expanded Materials and Methods in the Supplemental Material. All raw data and analytical methods are available from the corresponding authors on reasonable request. Sequencing data have been made publicly available at the Gene Expression Omnibus. The mouse scRNA-seq data can be accessed at GSEXXXX, and the mouse scATAC-seq data can be accessed at GSEXXXX. The mouse spatial transcriptomic data and computer code used in this study are available on request.

### Murine Studies

For experiments designed to visualize specific cellular markers, *Sting* flox/flox mice (strain No.035692), *Irf3* flox/flox mice (strain No.036260) and *Lyz2*-CreERT2 mice (strain No.032291) were obtained from The Jackson Laboratory and cross-bred. Floxed pups with a *Lyz2*-CreERT2+ genotype were given tamoxifen to induce the expression of Cre recombinase. This led to the *Sting* or *Irf3* knocking out in the MΦs. At 8 weeks of age, these mice underwent infusion with either saline or Ang II (Bachem, 4474-91-3, 1000ng/kg·min) for 7 days by osmotic pump (Alzet, model #2001), and aortas were collected at the 7-day time point for further analysis.

*Sting* knock out mice (strain No.025805) were purchased from The Jackson Laboratory, and male wildtype (WT) mice (C57BL/6J) were obtained from the Baylor College of Medicine animal facility. At 8 weeks of age, mice underwent osmotic pump implantation and were infused with either saline or Ang II (1000ng/kg·min) for 7 days. At the end of the infusion period, mice were euthanized by CO_2_ asphyxiation, followed by perfusion with 10 mL of ice-cold PBS through the left ventricle. The definition of aortic dissection is the same as those previously described^19^. For mice selected for scRNA-seq and scATAC-seq analysis, ascending aortic tissues were collected after the infusion period, and cells were prepared for single-cell suspension according to the protocol described previously, an additional step for single-nuclear obtain is necessary for scATAC-seq analysis as we described previously^20^. For mice selected for spatial transcriptomic analysis, ascending aortic tissues were fixed with 4% paraformaldehyde and processed for paraffin embedding. Then, within a base mold (10 × 10 × 5 mm) filled with melted paraffin, 9 aortas (3 control, 3 non-dissected, and 3 dissected) were arranged in a 3×3 array, before the paraffin was allowed to solidify.

### Human Tissue Study

Ascending aortic tissues from patients with sporadic ascending thoracic aortic aneurysm without dissection (ATAA, n=9; 6 men and 3 women), patients with acute ascending thoracic aortic dissection (ATAD, n=9; 6 men and 3 women), and donors of heart or lung transplants without aortic diseases (controls, n=8; 5 men and 3 women) were collected for scRNA-seq study. Aortic disease related to heritable conditions, infection, aortitis, or trauma were excluded from the study. The protocol for collecting human tissue samples was approved by the institutional review board at Baylor College of Medicine (Houston, TX). Written informed consent for participation was provided by patients or family members whenever possible and by organ donors’ legal representatives. For patients who were too ill to provide consent and for whom family members were not available, the IRB approved a waiver of consent. All experiments conducted with human tissue samples were performed in accordance with the relevant guidelines and regulations.

### Statistical Analysis

All data were analyzed with either GraphPad Prism V10.3.1 statistical analysis software (Graphpad Software, Boston, MA) or R V4.4.2. Quantitative data are presented as mean ± SD or as the median and interquartile range, as appropriate. Continuous variables were first assessed for normality using the Shapiro-Wilk normality test. Data not exhibiting a normal distribution were log2-transformed and retested for normality. To examine whether a particular class of genes had increased (or decreased) expression in RNA-seq data sets, the Wilcoxon Rank Sum test was performed to compare the genes between 2 groups; P values were adjusted by Bonferroni correction using all genes in the data set. Statistical analysis of quantitative polymerase chain reaction and chromatin immunoprecipitation assay data was performed using 2-way ANOVA, followed by the Dunnet multiple comparisons test when appropriate. For comparisons between two conditions, the choice of test was dependent on the data distribution. Equality of variance was examined by an F test. A two-tailed Student’s t test was used for normally distributed data with equivalent variance. For skewed data that remained non-normal after transformation, Mann-Whitney U test was used. Differences with P<0.05 were considered statistically significant.

## RESULTS

### Marked Pro-Inflammatory MΦ Phenotypes in Human ATAD

To delineate aortic cell dynamics during ATAD progression, we performed scRNA-seq of ascending aortic tissues from organ-donor controls without aortic diseases, patients with ATAA without dissection, and patients with ATAD (Figure 1A). In total, 27 human samples were analyzed.

**Figure 1.**
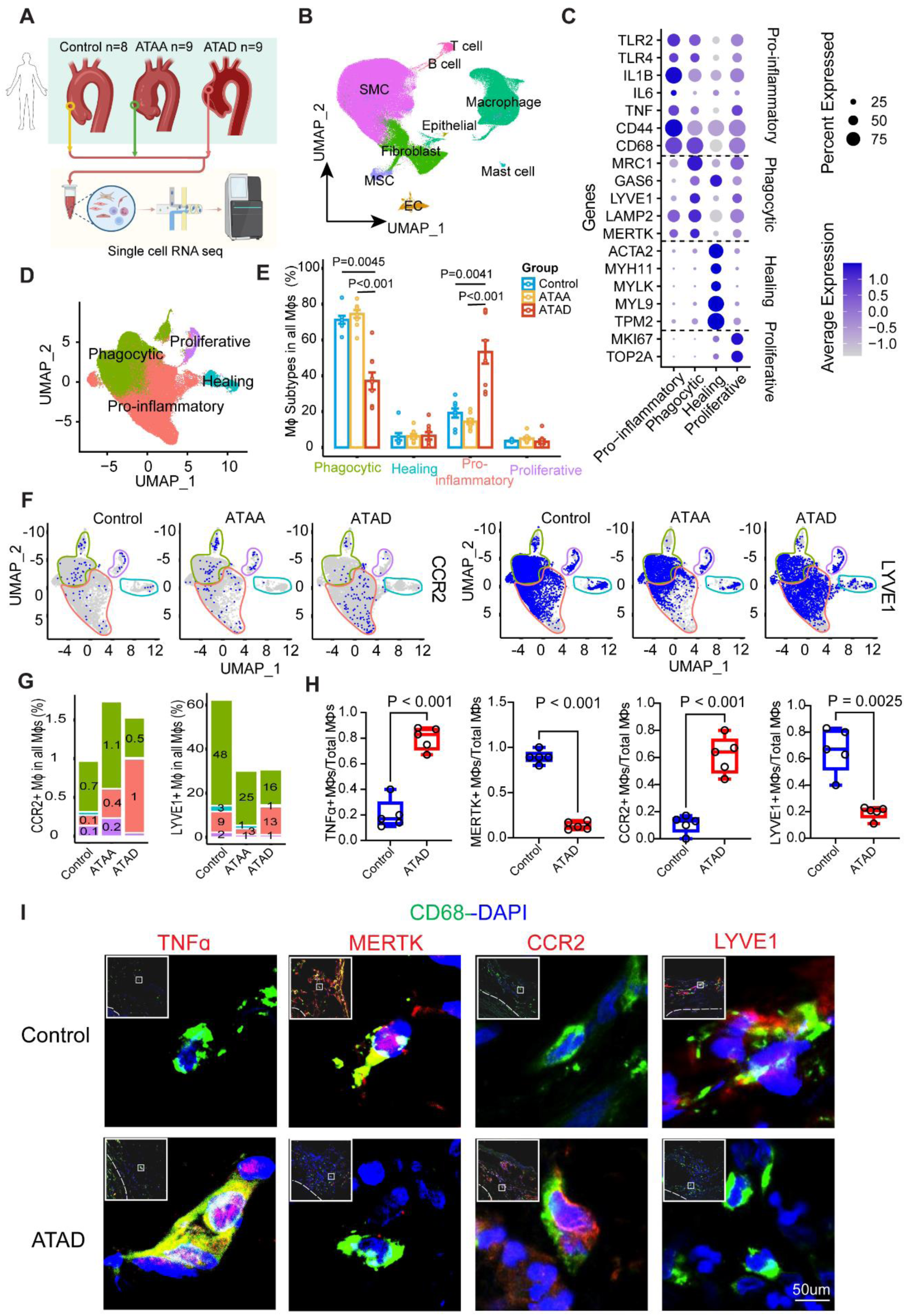
Diverse Phenotypes of MΦs Involved in Human Ascending Aortic Thoracic Dissection. **A,** Single-cell RNA sequencing (scRNA-seq) of human ascending aortic tissues from individuals without aortic disease (control), patients with ATAA, and patients with ATAD. **B,** Uniform manifold approximation and projection (UMAP) showing major cell types, including endothelial cells (ECs), smooth muscle cells (SMCs), fibroblasts (FBs), macrophages (MΦs), T lymphocytes, B lymphocytes, mast cells, epithelial cells and mesenchymal stem cells (MSCs). **C,** Dot plot showing enriched genes in each MΦ subclusters. **D,** UMAP showing MΦ subclusters with their annotations. **E,** Proportion of MΦ subclusters in all MΦs among groups. **F,** Dynamic UMAPs of *CCR2* and *LYVE1* positive cells among groups. **G,** Proportion of *CCR2* and *LYVE1* positive cells in all MΦs among groups. **H,** Statistical analysis of marker protein-positive MΦs/total MΦs in the same area size of IF images between control and ATAD groups. **I,** Immunofluorescence (IF) staining on human ascending aortic tissues. CD68 is the marker for MΦs, TNFα is the marker for pro-inflammatory macrophages, MERTK is the marker for phagocytic macrophages, CCR2 is the marker for recruited MΦs, and LYVE1 is the marker for resident MΦs.

Unsupervised clustering identified the major cellular populations within the aortic wall, with MΦs representing the predominant immune cell population (Figure 1B). Given their abundance, we focused subsequent analyses on MΦs and resolved several transcriptionally distinct subpopulations. Based on cluster-specific gene enrichment (Figure 1C), MΦs were classified as pro-inflammatory (*TLR2, TLR4, IL1B, IL6, TNF, CD44, CD68*), phagocytic (*MRC1, GAS6, LYVE1, LAMP2, MERTK*), healing/smooth muscle–like (*ACTA2, MYH11, MYLK, MYL9, TPM2*), and proliferative (*MKI67, TOP2A*) subclusters (Figure 1D).

Macrophage proportional distribution analysis revealed that overall MΦ composition in ATAA closely resembled that of control aortas. In contrast, ATAD samples exhibited a pronounced shift in MΦ composition, characterized by a marked expansion of pro-inflammatory MΦs and a concomitant reduction in phagocytic MΦs compared with controls (Figure 1E).

### Origin-Specific MΦ Reprogramming in Human ATAA and ATAD

We next examined MΦ origins using CCR2 as a marker of recruited MΦs^21^ and LYVE1 as a marker of tissue-resident MΦs^22^. *CCR2*⁺ MΦs were significantly increased in both ATAA and ATAD relative to controls (Figures 1F and 1G). Notably, in ATAA, CCR2⁺ MΦs were predominantly associated with phagocytic and pro-healing subpopulations, whereas in ATAD they were preferentially enriched within the pro-inflammatory MΦ cluster (Figures 1F and 1G). These findings suggest that recruited MΦs are selectively skewed toward a pro-inflammatory fate in ATAD.

In parallel, *LYVE1*⁺ resident MΦs were reduced in diseased aortas (Figures 1F and 1G). In ATAA, *LYVE1*⁺ MΦs primarily localized to phagocytic subclusters, whereas in ATAD they shifted toward pro-inflammatory states, indicating phenotypic reprogramming of resident MΦs from anti-inflammatory, phagocytic phenotypes to pro-inflammatory phenotypes during dissection.

Consistent with these transcriptomic findings, immunostaining demonstrated significant upregulation of the pro-inflammatory and recruited MΦ markers TNFα and CCR2 in ATAD aortas compared with controls, accompanied by marked reductions in the phagocytic and resident MΦ markers MERTK and LYVE1 (Figures 1H and 1I).

Collectively, these human data demonstrate that whereas ATAA largely preserves a MΦ landscape similar to controls, ATAD is characterized by a profound shift toward pro-inflammatory MΦ dominance with loss of phagocytic populations. Both recruited and resident MΦs contribute to this inflammatory bias, with *CCR2*⁺ recruited MΦs and *LYVE1*⁺ resident MΦs preferentially adopting pro-inflammatory phenotypes during ATAD progression.

### Pro-Inflammatory MΦ Modulation in Ang II–Induced ATAD

To investigate the aortic MΦ phenotypic modulation during ATAD development, we performed scRNA-seq analysis of the ascending aortic tissues from Ang II-induced ATAD mice. WT mice were given saline or Ang II for 7 days (Figure 2A).

**Figure 2.**
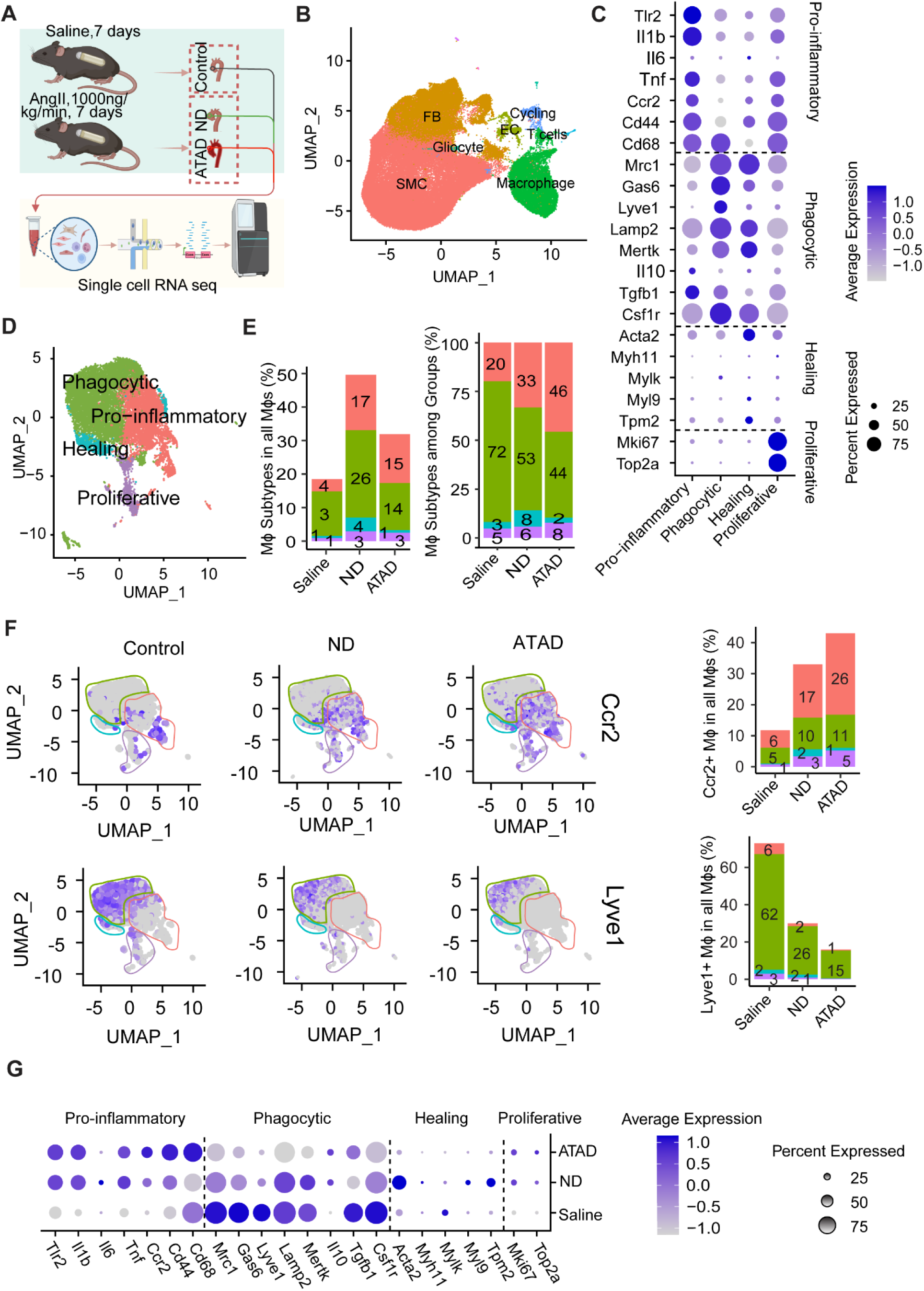
Pro-inflammatory Modulation of MΦs Involved in Mouse Ascending Aortic Thoracic Dissection. **A,** scRNA-seq of mouse ascending aortic tissues from individuals were infused with saline or Ang II with (ATAD) or without aortic dissection (ND). **B,** UMAP of major cell types, including EC, SMC, FB, MΦ, T cell, Cycling and Gliocyte. **C,** Dot plot showing enriched genes in each MΦ subclusters. **D,** UMAP showing MΦ subclusters with their annotations and proportion of MΦ subclusters in all MΦs and among groups **(E)**. **F,** Dynamic UMAPs and proportion of *Ccr2* and *Lyve1* positive cells among groups. **G,** Dot plot showing enriched genes in each MΦ subclusters among groups.

Major cell types were identified, and again MΦs showed the highest proportion among immune cells (Figure 2B). Based on cluster-specific gene enrichment (Figure 2C), the MΦ subpopulations (Figure 2D) were defined as pro-inflammatory (*Tlr2*, *Il1b*, *Il6*, *Tnf*, *Ccr2*, *Cd44*, and *Cd68*), phagocytic (*Mrc1*, *Gas6*, *Lyve1*, *Lamp2*, *Mertk*, *Il10*, *Tgfb1*, and *Csf1r*), healing (*Acta2*, *Myh11*, *Mylk*, *Myl9*, and *Tpm2*), and proliferative (*Mki67* and *Top2a*) subclusters (Figure 2D).

Comparative analysis of cell populations showed that MΦs were increased in both non-dissected (ND) and dissected (ATAD) groups (Figure 2E, left). However, this increase was accompanied by a progressive reduction in differentiation toward phagocytic MΦs and a concomitant expansion of pro-inflammatory MΦs (Figure 2E, right).

Consistent with the human dataset, Ang II infusion in mice resulted in a progressive increase in *Ccr2*⁺ recruited MΦs and a corresponding decrease in *Lyve1*⁺ resident MΦs (Figure 2F). Importantly, *Ccr2*⁺ recruited MΦs predominantly gave rise to pro-inflammatory MΦs, whereas *Lyve1*⁺ resident MΦs primarily contributed to phagocytic MΦ populations (Figure 2F).

Consistently, genes associated with pro-inflammatory responses were upregulated, while phagocytic gene programs were suppressed following Ang II infusion (Figure 2G). Together, these findings demonstrate that macrophage phenotypic remodeling in the mouse ATAD model closely mirrors the patterns observed in human aortic tissues.

### Spatial Reorganization of MΦ Subsets During Aortic Wall Injury

We next interrogated the spatial organization of MΦs and their subpopulations in relation to aortic injury and remodeling using spatial transcriptomic analysis of ascending aortic segments from Ang II– or saline-infused mice (Figure 3A). Integration with H&E-stained sections identified MΦs as a major immune population predominantly localized to the adventitial layer (Figure 3B). Spatial transcriptomics further resolved multiple MΦ subpopulations defined by distinct gene expression programs (Figures 3C and 3D).

**Figure 3.**
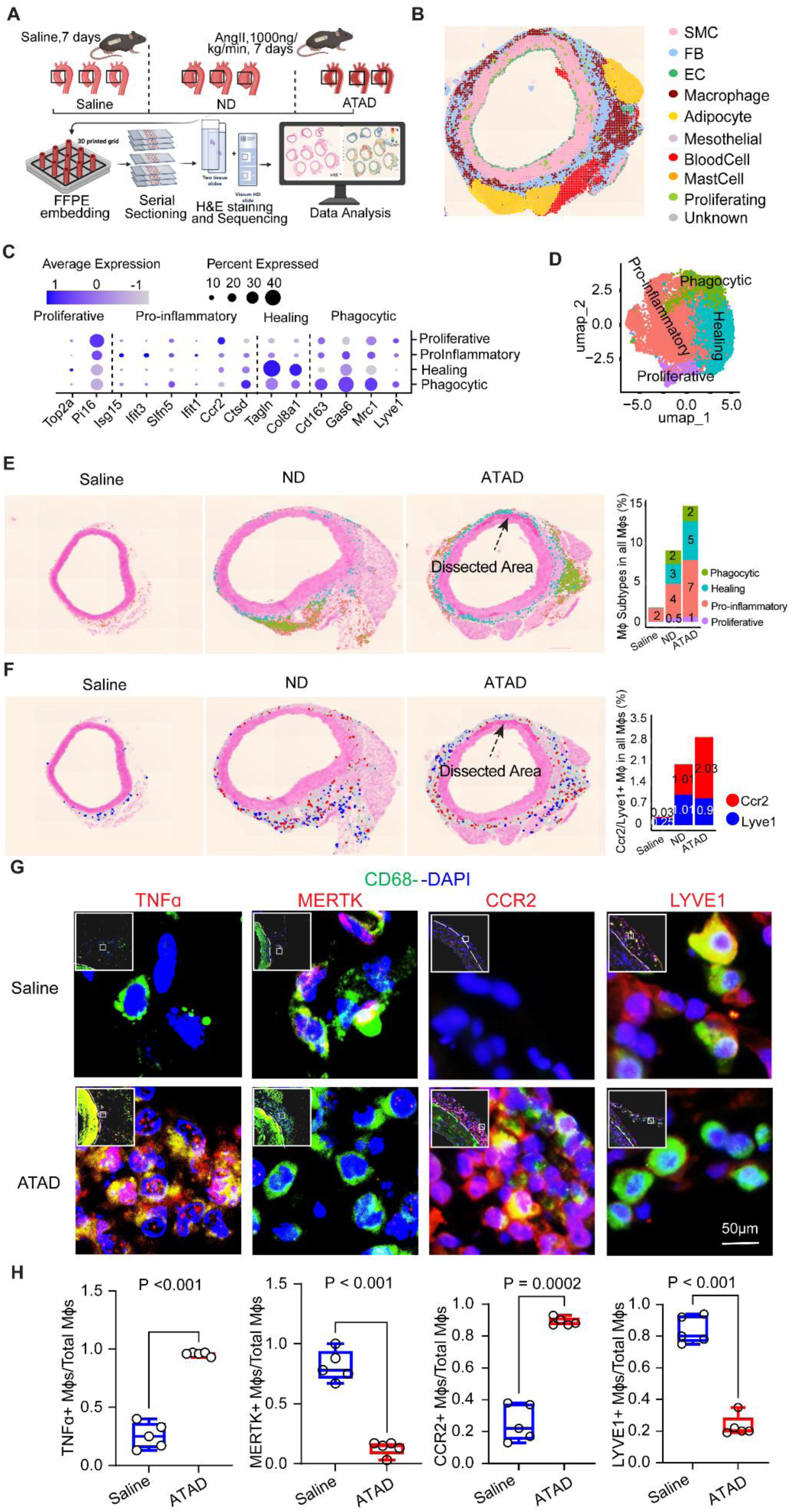
Phenotypic Modulated MΦs Were Recruited to the Dissected Area in Mouse Ascending Aorta. **A,** Spatial transcriptomic analysis of mouse ascending aortic tissues from individuals were infused with saline or Ang II with (ATAD) or without aortic dissection (ND). **B,** Major cell types identified from the spatial transcriptomic data were projected on a 2-dimensional plot. **C,** Dot plot showing enriched genes in each MΦ subclusters. **D,** UMAP showing MΦ subclusters with their annotations. **E,** Dynamic UMAPs and proportions of MΦ subclusters among groups. **F,** Dynamic UMAPs and proportions of *Ccr2* and *Lyve1* positive cells among groups. **G,** IF staining of mouse ascending aortic tissues. **H,** Statistical analysis of marker protein-positive MΦs/total MΦs in the same area size of IF images between saline and ATAD groups.

Notably, total MΦ abundance increased progressively from control to non-dissected (ND) and dissected (ATAD) groups (Figure 3E). This accumulation was spatially patterned: pro-inflammatory MΦs were enriched at sites of aortic dissection and tissue disruption, whereas phagocytic MΦs were concentrated within adventitial niche regions (Figure 3E). The spatial findings suggest that pro-inflammatory MΦs may positioned to actively drive local tissue damage and propagation of dissection, while phagocytic MΦs may contribute to debris clearance and tissue containment. Additionally, both *Ccr2*⁺ recruited and *Lyve1*⁺ resident MΦs expanded with disease progression and were present within both dissected regions and adventitial niches (Figure 3F).

Consistently, immunostaining demonstrated a marked increase in recruited MΦs (CCR2⁺CD68⁺) and pro-inflammatory MΦs (TNFα⁺Cd68⁺) and accompanied by a loss of resident MΦs (LYVE1⁺CD68⁺) and phagocytic MΦs (MERTK⁺CD68⁺) in ATAD compared with saline-infused controls (Figures 3G and 3H).

Together, these spatially patterned phenotypes suggest that local aortic microenvironmental cues may reprogram MΦs toward pro-inflammatory states, thereby providing a mechanistic framework to investigate the upstream signals and epigenetic regulators that drive this process.

### Pro-Inflammatory MΦ Modulation in Ang II–Induced ATAD Is Epigenetically Regulated

We next investigated the mechanisms underlying MΦ phenotypic modulation during ATAD development. Given the central role of epigenetic regulation and chromatin remodeling in controlling gene expression, we performed scATAC-seq on mouse ascending aortic tissues following saline or Ang II infusion for 7 days (Figure 4A). MΦs were readily identified based on cluster-specific chromatin accessibility at the *Cd68* locus (Figure 4B).

**Figure 4.**
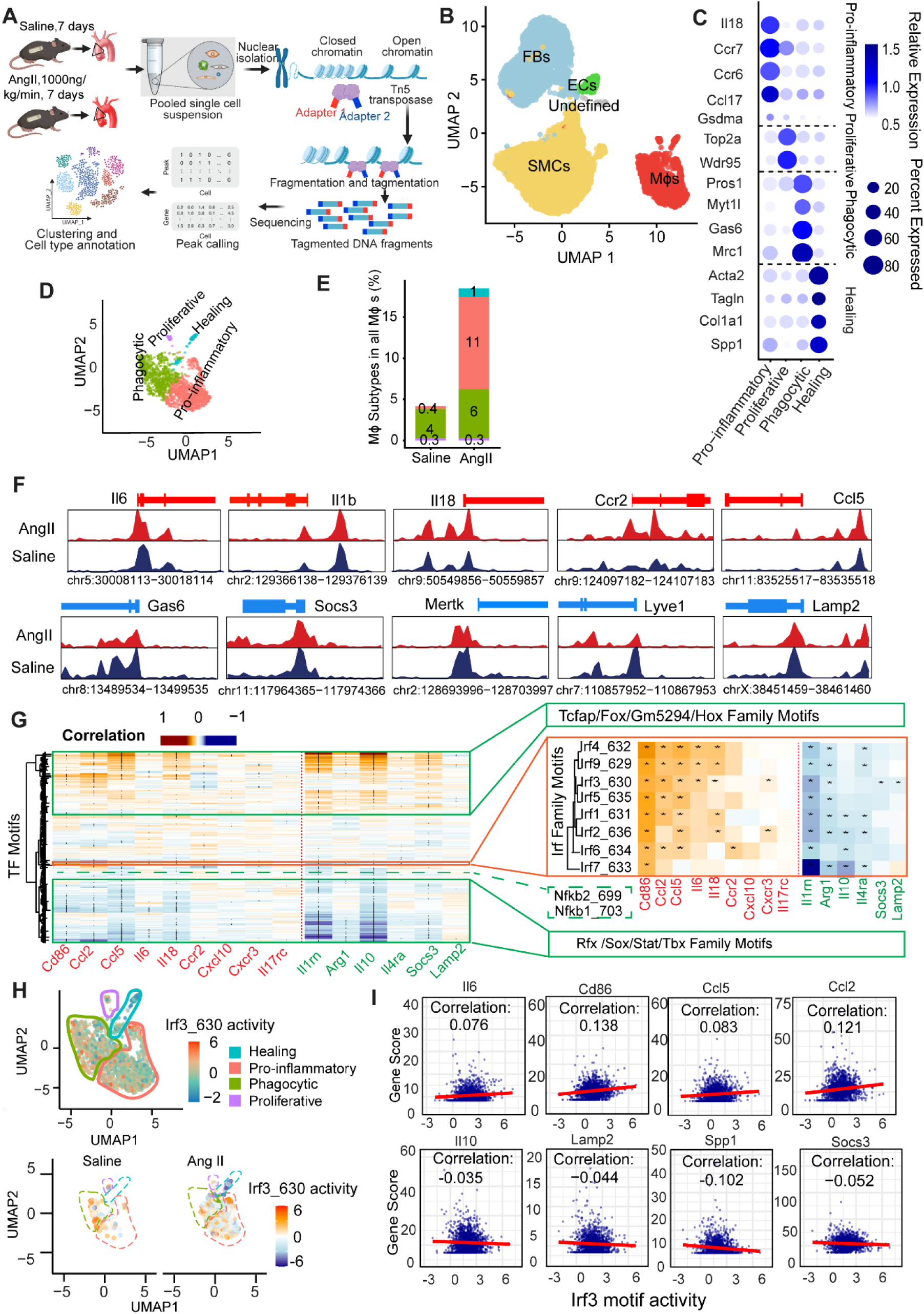
Ang II Infusion Increased Chromatin Accessibility in Pro-inflammatory Genes and Decreased Chromatin Accessibility in Phagocytic Genes. **A,** A schematic diagram illustrating the design for single-cell ATAC sequencing (scATAC-seq) studies in mice infused with saline or Ang II. **B,** Major cell types identified in scATAC-seq data from aortic tissues of saline and Ang II-infused mice. **C,** Dot plot showing enriched genes in each MΦ subcluster. **D,** UMAP showing MΦ subclusters with their annotations. **E,** Proportions of MΦ subclusters between groups. **F,** Chromatin accessibility at transcription starts sites of pro-inflammatory and phagocytic genes in saline or Ang II-infused groups. **G,** Heatmap showing correlation coefficients between chromatin accessibility of pro-inflammatory or phagocytic genes and motif activity of transcription factors in mouse aortic MΦs. **H,** Overall activity of the Irf3 motif in MΦs between saline and Ang II-infused groups. **I,** Dot plots showing the correlation of Irf3 motif activity with pro-inflammatory and phagocytic gene accessibility.

Based on distinct patterns of open chromatin at phenotype-defining genes, MΦs were further resolved into pro-inflammatory (*Il18, Ccr7, Ccr6, Ccl17,* and *Gsdma*), phagocytic (*Pros1, Myt1l, Gas6,* and *Mrc1*), healing (*Acta2, Tagln, Col1a1,* and *Spp1*), and proliferative (*Top2a* and *Wdr95*) subclusters (Figures 4C and 4D), indicating that MΦ phenotypic states are epigenetically encoded.

Importantly, Ang II infusion led to a marked expansion of MΦs compared with saline controls, with a preferential increase in pro-inflammatory MΦs (Figure 4E). Quantitative analysis further revealed an increased proportion of pro-inflammatory MΦs accompanied by a reduction in phagocytic MΦs, consistent with epigenetically driven phenotypic skewing toward a pro-inflammatory state.

Concordantly, Ang II infusion was associated with increased chromatin accessibility at transcription start sites of pro-inflammatory genes and decreased accessibility at phagocytic gene loci (Figure 4F). Together, these data demonstrate that Ang II promotes MΦ reprogramming through coordinated chromatin remodeling, providing direct epigenetic evidence for phenotypic polarization toward pro-inflammatory MΦs during ATAD development.

### IRF3 Is a Potential Epigenetic Driver of Pro-Inflammatory Programs While Suppressing Phagocytic Gene Programs

To identify transcription factors (TFs) that drive MΦ pro-inflammatory phenotype, we performed a correlation analysis linking motif activity/motif accessibility of >300 TFs with gene activity/chromatin accessibility of genes associated with pro-inflammatory or phagocytic MΦ phenotypes (Figure 4G). This analysis revealed three distinct regulatory patterns.

In the first group, motif activities of TFs (e.g., Tcfap, Fox, Gm5294, and Hox families) positively correlated with both pro-inflammatory and anti-inflammatory/phagocytic gene programs, suggesting shared roles in broadly activating MΦ transcriptional responses. Notably, despite its well-established role in inflammatory signaling^23,24^, NF-κB motif activity did not significantly correlate with either gene program, indicating that NF-κB–dependent signaling alone may be insufficient to explain the observed chromatin remodeling.

In contrast, in the second group, motif activities of TFs (e.g., Rfx, Sox, Stat, and Tbx family) showed negative correlations with both pro-inflammatory and anti-inflammatory/phagocytic gene programs, consistent with a potential inhibitory role in overall MΦ activation.

Notably, the third transcription factor group, dominated by the Irf family, exhibited a distinctive regulatory pattern in which motif activity was positively correlated with pro-inflammatory gene activity and inversely correlated with anti-inflammatory/phagocytic programs (Figure 4G). This pattern implicates IRF TFs as key drivers of MΦ polarization, simultaneously promoting inflammatory gene programs while repressing anti-inflammatory/phagocytic genes.

Among these, Irf3 emerged as a leading regulator. Irf3 motif activity was increased in MΦs, particularly in the pro-inflammatory MΦs, in Ang II–infused aortas compared with saline controls (Figure 4H). Additionally, Irf3 motif activity positively correlated with the activity of key pro-inflammatory genes (e.g., *Il6, Cd86, Ccl5,* and *Ccl2*), while inversely correlated with anti-inflammatory/phagocytic genes (e.g., *Il10, Lamp2, Spp1,* and *Socs3*) (Figure 4I). Collectively, these findings identify IRF3 as a central transcriptional regulator that promotes pro-inflammatory macrophage programming while suppressing anti-inflammatory/phagocytic gene networks in response to Ang II stimulation.

### STING–IRF3 Drives Epigenetic Reprogramming of MΦs Toward Pro-Inflammatory Phenotypes

IRF3 is a key pro-inflammatory TF activated by DNA damage and cytosolic DNA sensing through the cGAS–cGAMP–STING pathway^25,26^. We have previously demonstrated activation of cytosolic DNA–STING signaling and its importance in ATAD development^27^. Building on these observations, we investigated whether STING–IRF3 signaling directly regulates inflammatory gene programs and MΦ phenotypes.

To this end, THP-1 cells were transfected with STING or IRF3 siRNAs followed by cGAMP stimulation (Figure 5A), with efficient knockdown confirmed by immunoblotting (Figure 5B). Bulk RNA-seq revealed distinct transcriptional profiles among experimental groups (Figure 5C). cGAMP stimulation robustly induced pro-inflammatory gene expression while suppressing phagocytic gene expression (Figures 5D and 5E). Importantly, silencing either STING or IRF3 abrogated these transcriptional changes, a finding further validated by RT-PCR (Figure 5F), establishing a direct role for the STING–IRF3 axis in promoting inflammatory gene expression while repressing phagocytic programs.

**Figure 5.**
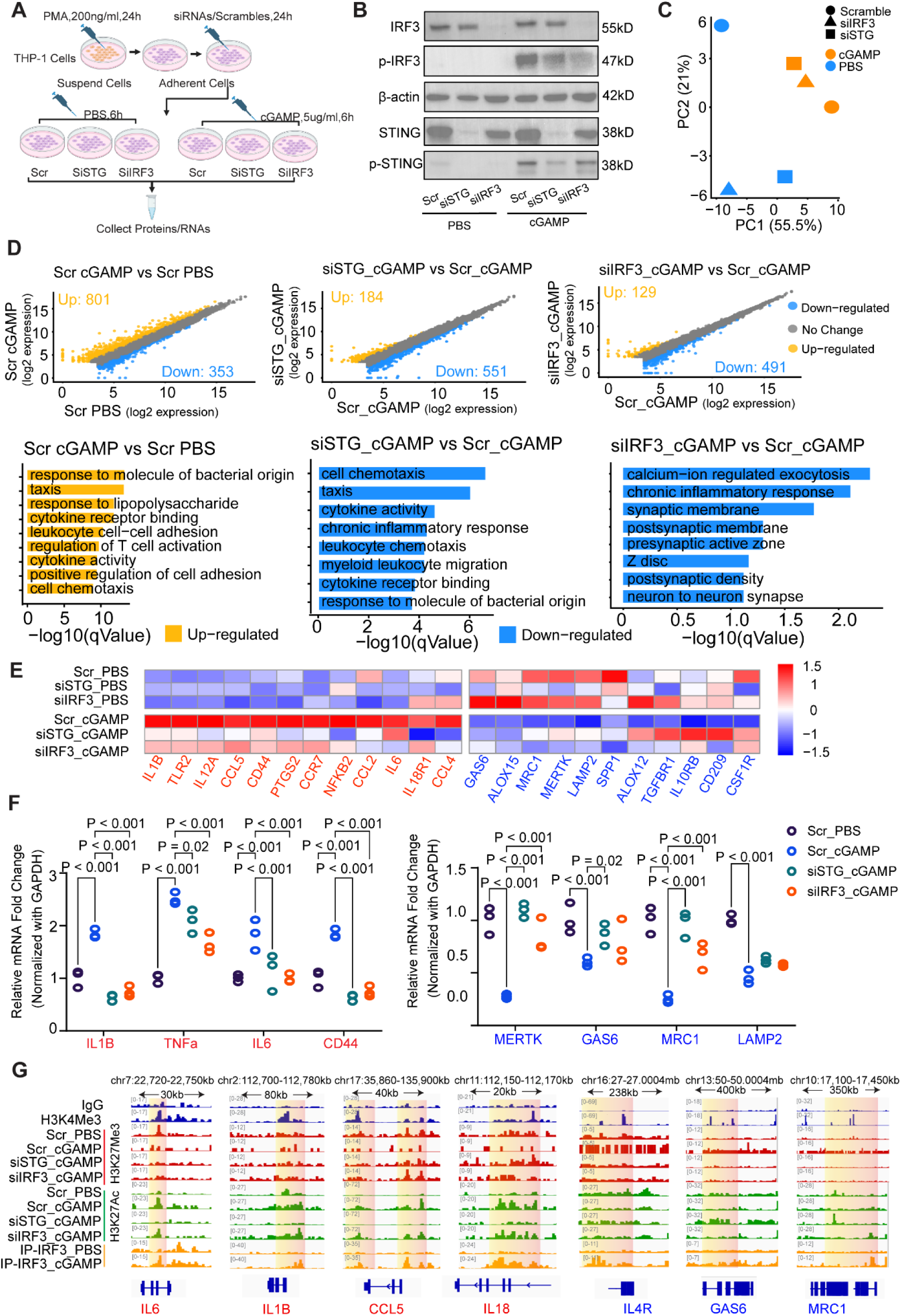
IRF3 as a Potential Driver for the Open Chromatin of Pro-inflammatory Genes and Suppression of Phagocytic Genes. **A,** A schematic diagram illustrating the design for the in vitro study. **B,** Western blotting showing the expression levels of IRF3, phosphorylated IRF3, STING, and phosphorylated STING following siRNA and cGAMP administration. **C,** PCA plot of bulk RNA sequencing among groups. **D,** Scale plots showing differentially expressed genes (DEGs) between groups, along with gene ontology (GO) analysis illustrating the functions of DEGs. **E,** Heatmap showing the expression of pro-inflammatory and phagocytic genes among groups. **F,** RT-PCR results of pro-inflammatory and phagocytic genes among groups. **G,** CUT&RUN sequencing showing the H3K27me3 and H3K27ac levels of pro-inflammatory and phagocytic genes among groups, including overlapped peaks of IRF3 binding sites and chromatin remodeling sites of genes.

We next examined whether STING–IRF3 signaling mediates these effects through chromatin remodeling. CUT&RUN-seq profiling of histone marks revealed that cGAMP stimulation decreased the repressive marker H3K27me3 and increased the active marker H3K27ac at pro-inflammatory gene loci, consistent with permissive chromatin remodeling (Figure 5G). Conversely, cGAMP increased H3K27me3 and reduced H3K27ac at phagocytic gene loci, indicating transcriptionally repressive chromatin remodeling. Notably, these chromatin changes were reversed by knockdown of either STING or IRF3, demonstrating that STING–IRF3 signaling might be required for epigenetic regulation of these gene programs.

To determine whether IRF3 directly engages these regulatory regions, we performed IRF3 CUT&RUN-seq. IRF3 binding was induced by cGAMP stimulation and localized to regulatory elements of both pro-inflammatory and phagocytic genes, overlapping with sites of chromatin remodeling (Figure 5G). These findings indicate that IRF3 may directly coordinate chromatin state and transcriptional output at target loci.

Collectively, these results demonstrate that activation of the STING–IRF3 pathway epigenetically programs MΦs toward a pro-inflammatory phenotype by opening chromatin at pro-inflammatory genes while enforcing repressive chromatin states at anti-inflammatory and phagocytic gene loci.

### IRF3 Induces Pro-Inflammatory Gene Expression via BRG1-Mediated Open Chromatin Remodeling

To further define the epigenetic mechanisms by which IRF3 induces pro-inflammatory gene activation while suppressing phagocytic programs in MΦs, we sought to identify chromatin/histone modifiers that interact with IRF3. We performed immunoprecipitation followed by mass spectrometry (IP–MS) in IRF3-overexpressing cells (Figure 6A). This analysis identified 665 IRF3-interacting proteins (Figure 6B) including proteins annotated with ATP-dependent activity acting on DNA (Figure 6C). Among these proteins, five histone modifiers/chromatin remodelers were identified including RUVBL1, RUVBL2, CHD4, SMARCA5, and SMARCA4 (BRG1) (Figure 6C).

**Figure 6.**
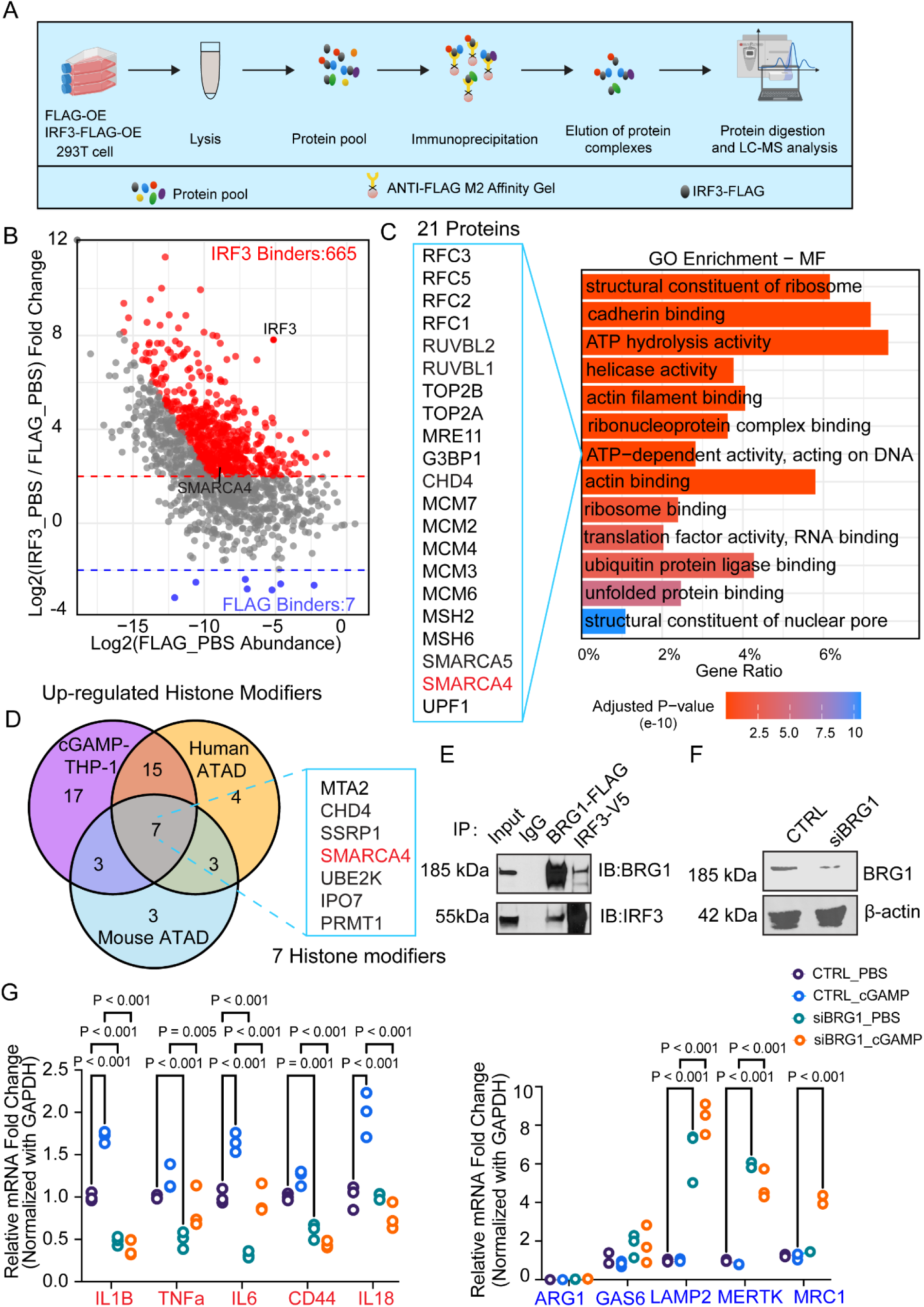
Promotion of Pro-inflammatory Genes and Suppression of Phagocytic Genes Expression by IRF3 Through the Recruitment of BRG1. **A,** Schematic diagram illustrating the design for the IP-MS study. **B,** Scatter plot of IP-MS showing the IRF3-specific binders and non-specific binders. **C,** GO enrichment of molecular functions of IRF3-specific binders. **D,** Venn plot of up-regulated histone modifiers in ATAD tissues of human and mouse scRNA-seq datasets and bulk RNA-seq dataset of THP1 cells within cGAMP stimulation. **E,** CO-IP results of BRG1-FLAG and IRF3-V5 interactions. **F,** Western blotting showing the expression of BRG1 following siBRG1 administration. **G,** RT-PCR results of pro-inflammatory and phagocytic genes among groups.

To select functionally relevant candidates, we integrated expression profiles of these IRF3-interacting histone modifiers across MΦs in human and mouse scRNA-seq datasets, together with bulk RNA-seq data from THP-1 cells. SMARCA4/BRG1 emerged as the most highly expressed candidate in MΦs from ATAD tissues and cGAMP administrated THP-1 cells (Figure 6D and Figure S1). Co-immunoprecipitation confirmed a physical interaction between IRF3 and BRG1 (Figure 6E), and immunostaining demonstrated nuclear co-localization of IRF3 and BRG1 following cGAMP stimulation (Figure S2).

We next examined the functional role of BRG1 in regulating MΦ gene expression. BRG1 is the catalytic ATPase subunit of the SWI/SNF chromatin-remodeling complex, which promotes chromatin accessibility and transcriptional activation, and has been implicated in inflammatory gene regulation in immune cells^28^. Consistent with this role, siRNA-mediated knockdown of BRG1 (Figure 6F) significantly suppressed pro-inflammatory gene expression under both basal conditions and following cGAMP stimulation (Figure 6G). Notably, BRG1 depletion also increased the expression of phagocytic genes, suggesting that BRG1 simultaneously promotes inflammatory programs while restraining reparative MΦ functions, a dual regulatory role that warrants further mechanistic investigation.

Collectively, these data support a model in which IRF3 recruits the SWI/SNF ATPase BRG1 to pro-inflammatory gene loci, promoting permissive chromatin remodeling and transcriptional activation while indirectly repressing phagocytic gene programs, thereby driving MΦ polarization toward a pro-inflammatory state.

### Macrophage-Specific STING Drives Pro-Inflammatory MΦ Programming and ATAD Development

We next examined the role of STING in aortic MΦ phenotypic modulation using single-cell RNA-seq data from ascending aortas of global *Sting* knockout (*Sting⁻/⁻*) and WT mice infused with Ang II or saline for 7 days. Ang II infusion in WT mice markedly increased the pro-inflammatory MΦ population while concomitantly reducing phagocytic MΦ population; these effects were substantially attenuated in *Sting*⁻/⁻ mice (Figure 7A). Consistently, Ang II–induced upregulation of pro-inflammatory genes and suppression of phagocytic genes were significantly blunted in *Sting*⁻/⁻ mice (Figure 7B). Together, these findings indicate that STING deficiency mitigates Ang II–induced pro-inflammatory MΦ programming in the aorta.

**Figure 7.**
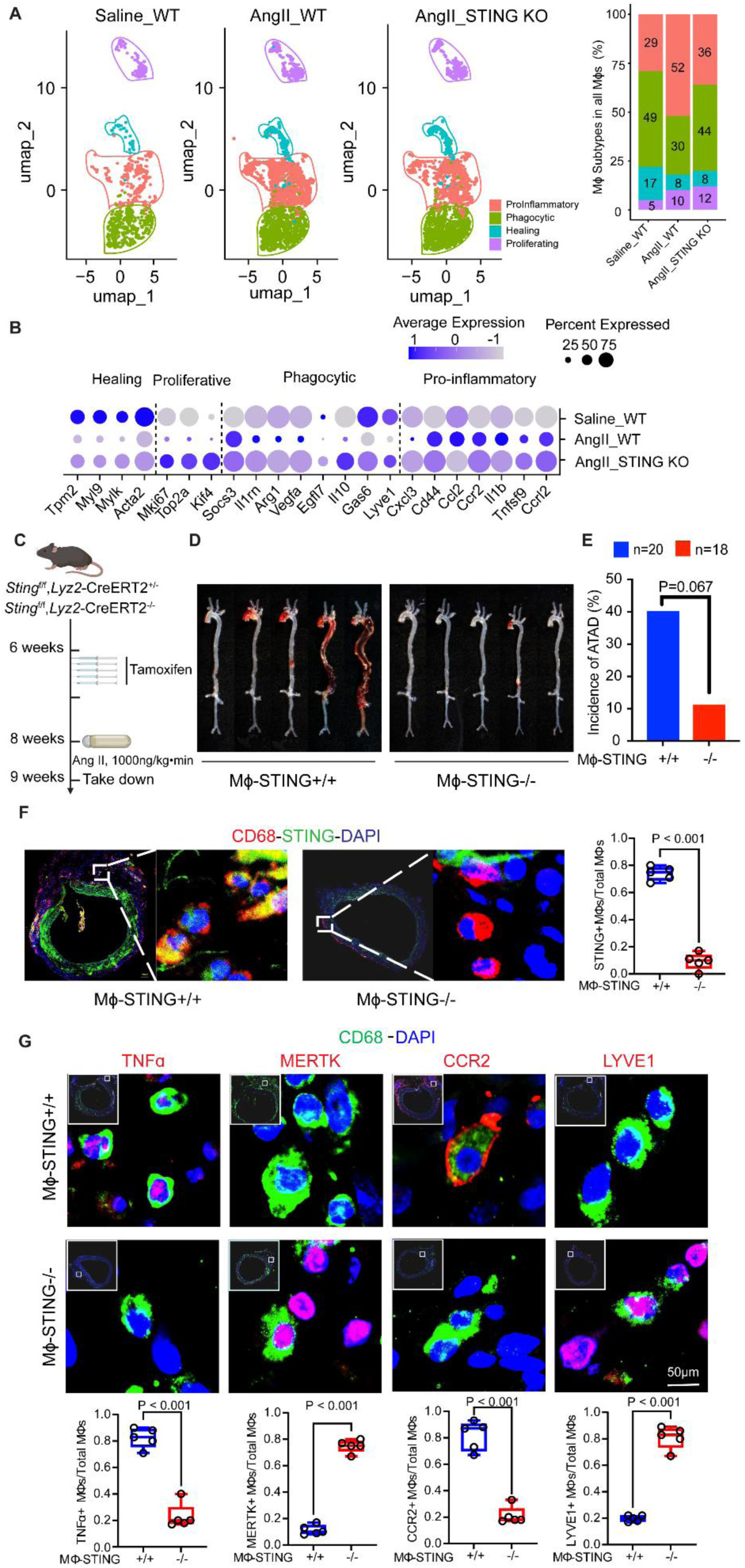
Global and MΦ-Specific *Sting* Deficiency Rescued Ang II-Induced Phenotypic Modulation of MΦ and ATAD Development in-vivo. **A,** Dynamic UMAPs and proportions of MΦ subclusters among groups. **B,** Dot plot showing enriched genes in each MΦ subclusters among groups. **C,** Workflow in MΦ-specific *Sting* deletion mice. **D,** Gross images of aortas in MΦ-STING⁺/⁺ and MΦ-STING⁻/⁻ mice. **E,** Incidence analysis of ATAD between MΦ-STING⁺/⁺ and MΦ-STING⁻/⁻ groups. **F,** IF staining of STING between MΦ-STING⁺/⁺ and MΦ-STING⁻/⁻ groups. **G,** IF staining and statistical analysis of MΦ marker proteins between MΦ-STING⁺/⁺ and MΦ-STING⁻/⁻ groups.

To define the contribution of MΦ-specific STING to ATAD development, we generated mice with MΦ specific Sting deletion (*Sting* flox/flox;*Lyz2*-Cre, MΦ-STING⁻/⁻) and used Cre-negative littermates (*Sting* flox/flox; MΦ-STING⁺/⁺) as controls. Following tamoxifen induction, mice were infused with Ang II for 7 days (Figure 7C). Compared with controls, MΦ-Sting⁻/⁻ mice exhibited significantly reduced aortic structural destruction (Figure 7D) and a lower incidence of ATAD (Figure 7C and Figure S3), supporting a critical role for MΦ -intrinsic STING in ATAD pathogenesis.

### Macrophage-Specific IRF3 Drives Ang II–Induced Pro-Inflammatory MΦ Reprogramming and ATAD Development

Finally, we examined the role of MΦ -specific IRF3 in MΦ phenotypic modulation and ATAD development. We generated mice with MΦ-restricted Irf3 deletion (*Irf3* flox/flox;*Lyz2*-Cre, MΦ-IRF3^⁻/⁻^) and used Cre-negative littermates (*Irf3* flox/flox; MΦ-IRF3⁺/⁺) as controls (Figure 8A). Following tamoxifen-induced Cre activation, these mice were given Ang II infusion for 7 days (Figure 8A).

**Figure 8.**
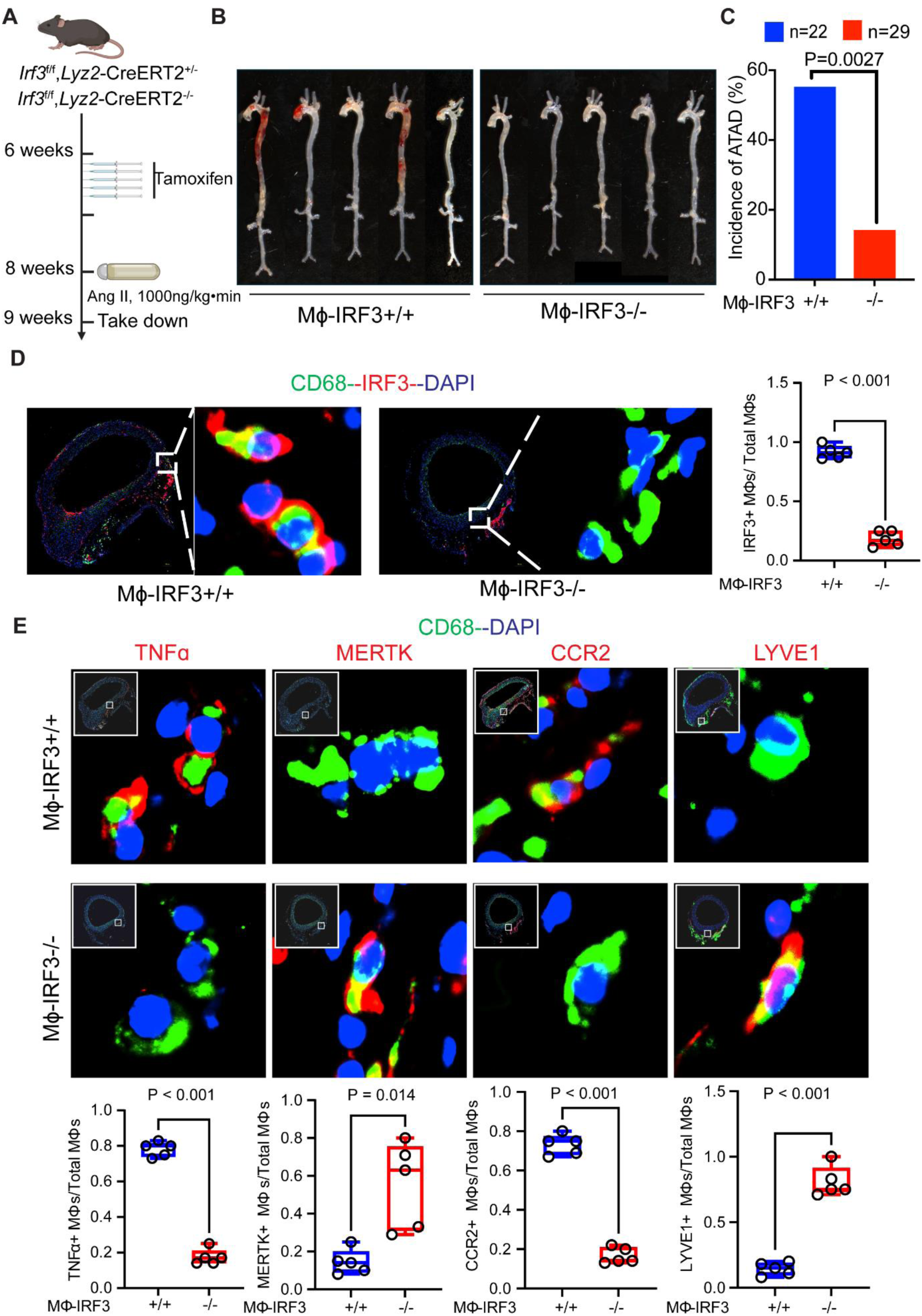
Macrophage-Specific *Irf3* Deficiency Reduced ATAD Development. **A,** Workflow in MΦ-specific *Irf3* deletion mice. **B,** Gross images of aortas in MΦ-IRF3⁺/⁺ and MΦ-IRF3⁻/⁻ mice. **C,** Incidence analysis of ATAD between MΦ-IRF3⁺/⁺ and MΦ-IRF3⁻/⁻ groups. **D,** IF staining of IRF3 between MΦ-IRF3⁺/⁺ and MΦ-IRF3⁻/⁻ groups. **E,** IF staining and statistical analysis of MΦ marker proteins between MΦ-IRF3⁺/⁺ and MΦ-IRF3⁻/⁻ groups.

Compared with control mice, MΦ-IRF3⁻/⁻ mice exhibited significantly reduced aortic structural destruction (Figure 8B) and a lower incidence of ATAD (Figure 8C and Figure S4). Immunostaining analyses demonstrated markedly decreased expression of STING, IRF3, TNFα, and CCR2, alongside increased expression of MERTK and LYVE1, in MΦ-IRF3⁻/⁻ mice compared with littermate controls (Figures 7D–E, and Figures 8D–E). Collectively, these findings support a critical role for MΦ-intrinsic IRF3 in driving pro-inflammatory MΦ reprogramming and promoting ATAD development.

## DISCUSSION

This study provides comprehensive evidence that aortic inflammation in sporadic ATAD is driven by epigenetically programmed MΦ reprogramming toward pro-inflammatory phenotypes. By integrating human single-cell transcriptomics, mouse single-cell multi-omics, spatial transcriptomics, and functional genetic models, we identified STING–IRF3–dependent epigenetic remodeling as a central mechanism that enforces inflammatory MΦ states while suppressing phagocytic and reparative programs. These findings offer a unifying mechanistic framework explaining the chronic, unresolved inflammation characteristic of ATAD and highlights MΦ epigenetic plasticity as a critical determinant of disease progression.

A major finding of this study is the clear divergence in MΦ composition between ATAA and ATAD. Despite increased MΦ accumulation in both conditions, ATAA largely preserved a balance of MΦ phenotypes similar to controls, with recruited *CCR2*⁺ MΦs retaining phagocytic and pro-healing features. In contrast, ATAD was marked by a pronounced expansion of pro-inflammatory MΦs accompanied by a loss of phagocytic populations. Importantly, both recruited and resident MΦs underwent phenotypic redirection in ATAD, indicating that inflammatory bias is not solely driven by monocyte-derived MΦ recruitment but also by local reprogramming within the aortic wall. These observations extend prior work describing inflammatory cell accumulation in aortic disease by demonstrating that phenotypic fate, rather than MΦ abundance alone, distinguishes dissection-prone aortas^27,29,30^. The convergence of recruited *CCR2*⁺ MΦs and resident *LYVE1*⁺ MΦs toward pro-inflammatory states in ATAD suggests that the diseased aortic microenvironment actively overrides intrinsic MΦ identity, promoting maladaptive inflammatory programs.

Our spatial transcriptomic and immunostaining analyses further revealed that MΦ phenotypes are not randomly distributed but are spatially organized in relation to aortic injury. Pro-inflammatory MΦs preferentially localize to regions of medial disruption and dissection, whereas phagocytic MΦs occupy adventitial niches. This spatial segregation supports a model in which pro-inflammatory MΦs actively contribute to local matrix degradation, smooth muscle cell loss, and propagation of dissection, while phagocytic MΦs may serve protective roles in debris clearance and tissue containment. The spatial coexistence of recruited and resident MΦs within both damaged and adventitial regions further supports the concept that local environmental cues, rather than MΦ origin alone, dictate functional polarization^31^. These findings underscore the importance of spatially resolved approaches in dissecting immune mechanisms in vascular disease.

A central advance of this work is the demonstration that MΦ phenotypic skewing in ATAD is encoded at the epigenetic level. Single-cell chromatin accessibility profiling revealed coordinated opening of pro-inflammatory gene loci and repression of phagocytic gene loci following Ang II stimulation. These chromatin changes closely mirrored transcriptional outputs and MΦ subtype composition^31^, providing direct evidence that MΦ fate decisions in ATAD are stabilized through epigenetic remodeling. This epigenetic framework offers an explanation for the persistence of inflammation in ATAD despite removal of acute triggers and aligns with emerging concepts of trained immunity. While trained immunity has been implicated in atherosclerosis and other chronic inflammatory conditions^14–16,32,33^, our study extends this paradigm to aortic disease and identifies MΦs within the aortic wall as key substrates for maladaptive immune training.

Through integrative motif analysis, we identified IRF3 as a unique transcriptional regulator that simultaneously promotes pro-inflammatory gene programs while suppressing phagocytic gene networks. Unlike canonical inflammatory transcription factors such as NF-κB^34^, IRF3 motif activity correlated bidirectionally with opposing MΦ programs^35^, suggesting a specialized role in enforcing phenotypic polarization rather than generalized activation. Mechanistically, we demonstrated that IRF3 directly binds regulatory regions of both pro-inflammatory and phagocytic genes and coordinates chromatin remodeling at these loci. These findings place IRF3 at the nexus of inflammatory signaling and epigenetic control, enabling sustained MΦ reprogramming beyond transient signaling events.

Our data further identified STING as the upstream activator of IRF3-mediated MΦ reprogramming in ATAD. Building on prior evidence of cytosolic DNA accumulation and STING activation in the diseased aorta^25–27^, we found that STING–IRF3 signaling is both necessary and sufficient to induce pro-inflammatory transcriptional and epigenetic changes in MΦs. A key mechanistic insight of this study is the identification of BRG1 as an IRF3-interacting chromatin remodeler that facilitates permissive chromatin states at inflammatory gene loci. IRF3 recruitment of BRG1 provides a direct molecular link between innate immune sensing and ATP-dependent chromatin remodeling. This mechanism explains how transient STING activation can produce durable changes in MΦ identity and function. While our data clearly demonstrated BRG1-dependent activation of pro-inflammatory genes, the mechanisms underlying IRF3-mediated repression of phagocytic programs remain incompletely defined. Preliminary evidence suggests involvement of repressive histone modifications, raising the possibility that IRF3 may coordinate both activating and repressive chromatin machinery to enforce phenotypic asymmetry. Future studies will be required to fully elucidate these repressive pathways.

The functional relevance of this epigenetic program is underscored by our genetic models. MΦ-specific deletion of *Sting* or *Irf3* markedly attenuated MΦ inflammatory reprogramming and reduced aortic destruction and ATAD incidence. These findings establish that STING–IRF3 signaling within MΦs is not merely correlative but is causally required for disease progression. Importantly, these effects were achieved without targeting other vascular cell types, highlighting MΦs as a critical therapeutic node. Given the essential roles of STING and IRF3 in host defense, MΦ-specific or context-dependent modulation of this pathway may offer a strategy to mitigate vascular inflammation while preserving systemic immunity.

Collectively, our findings identify epigenetic MΦ reprogramming as a fundamental driver of ATAD and position the STING–IRF3 axis as a promising therapeutic target. Pharmacologic modulation of trained immunity or chromatin remodeling—potentially through targeted nanotherapeutics or epigenetic inhibitors—may represent novel strategies to prevent disease progression in patients with ATAA at risk for dissection. Several limitations warrant consideration. Human samples represent late-stage disease and do not capture early initiating events. Additionally, while Ang II infusion recapitulates key inflammatory features of ATAD, it does not model all aspects of human disease. Finally, the reversibility of MΦ epigenetic programming remains to be determined and represents an important avenue for future investigation.

In summary, this study reveals that ATAD is driven by STING–IRF3–dependent epigenetic reprogramming of MΦs toward pro-inflammatory states. By integrating immune sensing, chromatin remodeling, and spatial context, our work provides a mechanistic blueprint for chronic aortic inflammation and identifies MΦ epigenetic plasticity as a therapeutic vulnerability in this devastating disease.

## ACKNOWLEDGEMENTS

The graphic illustration was created using BioRender.

## DISCLOURE OF INTEREST

Dr. LeMaire serves as a consultant for Cerus. Dr. Coselli serves as principal investigator, consults for, and receives royalties and a departmental educational grant from Terumo Aortic; consults and participates in clinical trials for Medtronic, Inc., and W.L. Gore & Associates; and participates in clinical trials for Abbott Laboratories, CytoSorbents, Edwards Lifesciences, and Artivion. The other authors report no conflicts.

---

**Figure S1.**
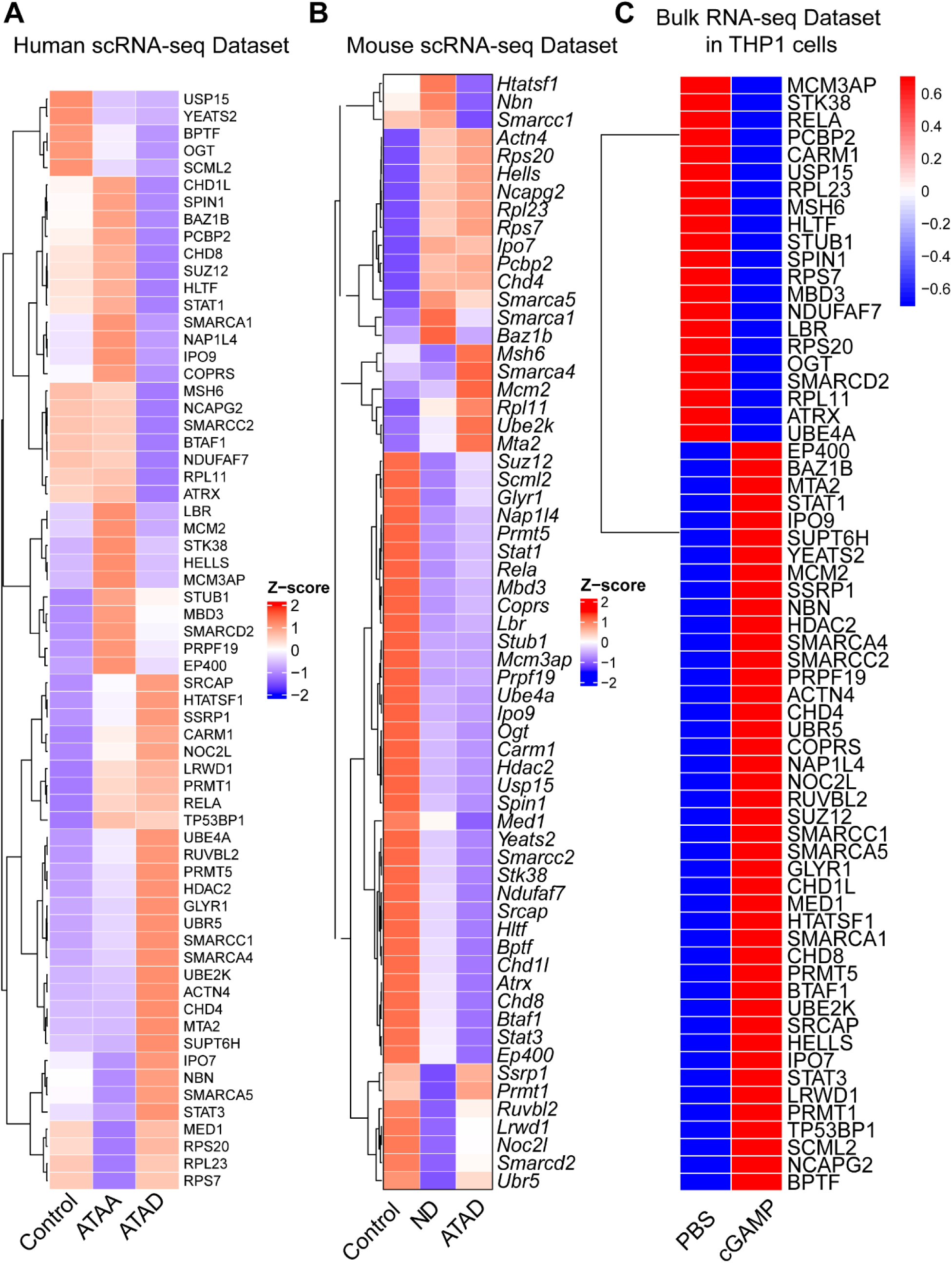
Heatmaps of Histone Modifiers in Different Datasets. **A.** Expression of histone modifiers in human scRNA-seq dataset, mouse scRNA-seq dataset **(B),** and in bulk RNA-seq dataset of THP-1 cells **(C).**

**Figure S2.**
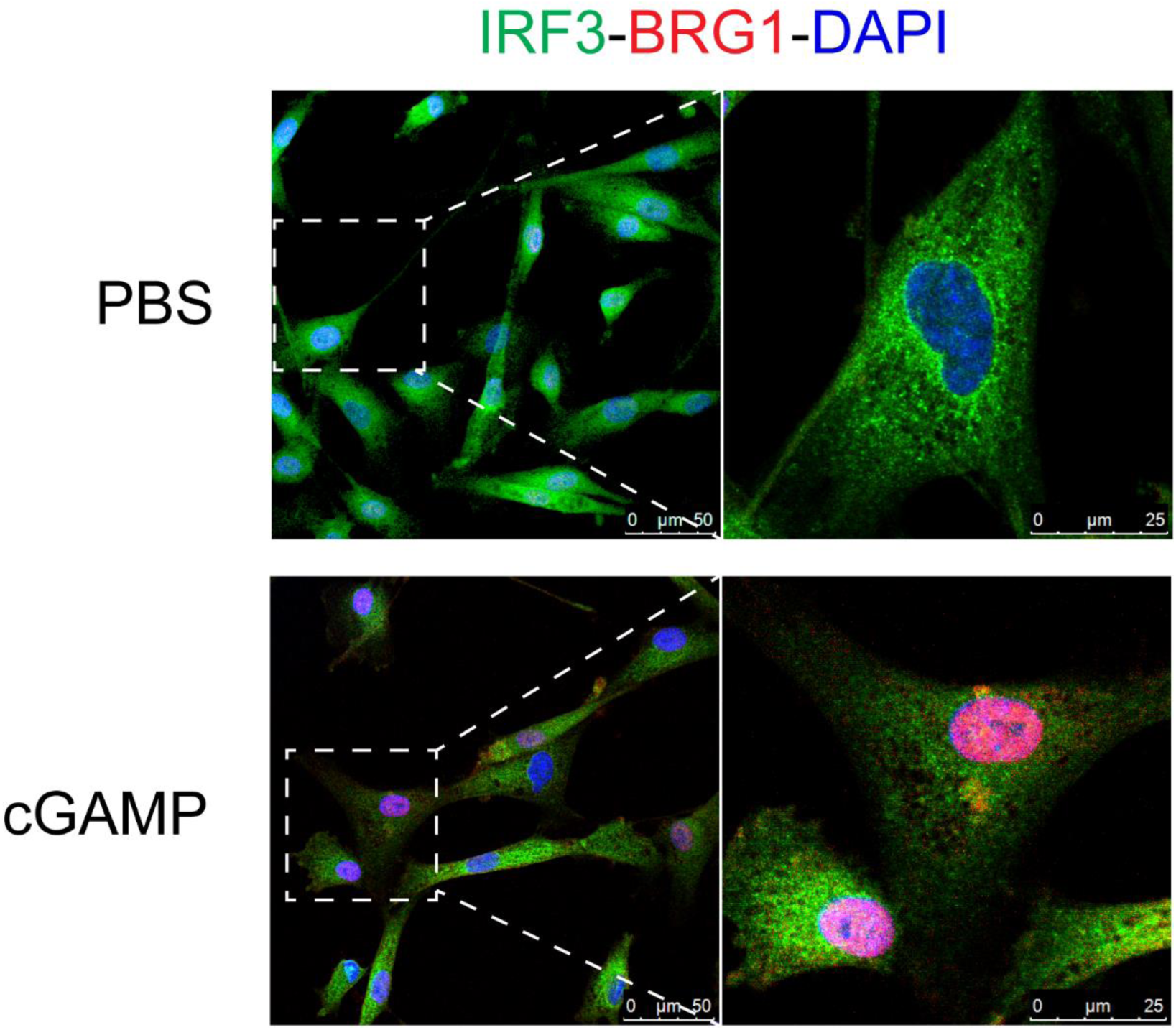
IF staining of the colocalization of the individual components of the IRF3-BRG1 complex in 293T cells.

**Figure S3.**
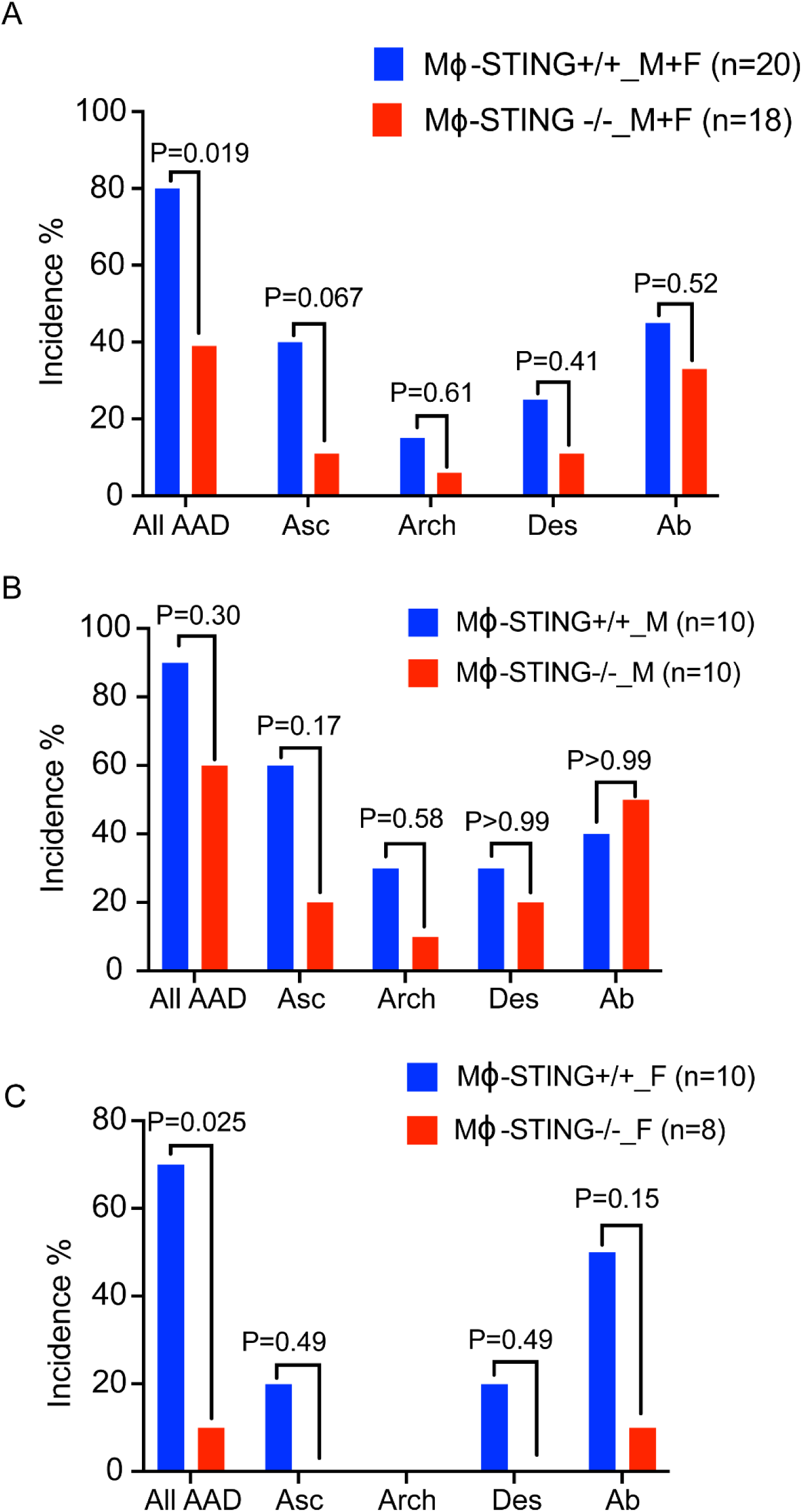
Incidence of aortic dissection in different aortic segments in all MΦ-specific *Sting* deficient mice **(A)**, and in male **(B)** and female **(C)** mice.

**Figure S4.**
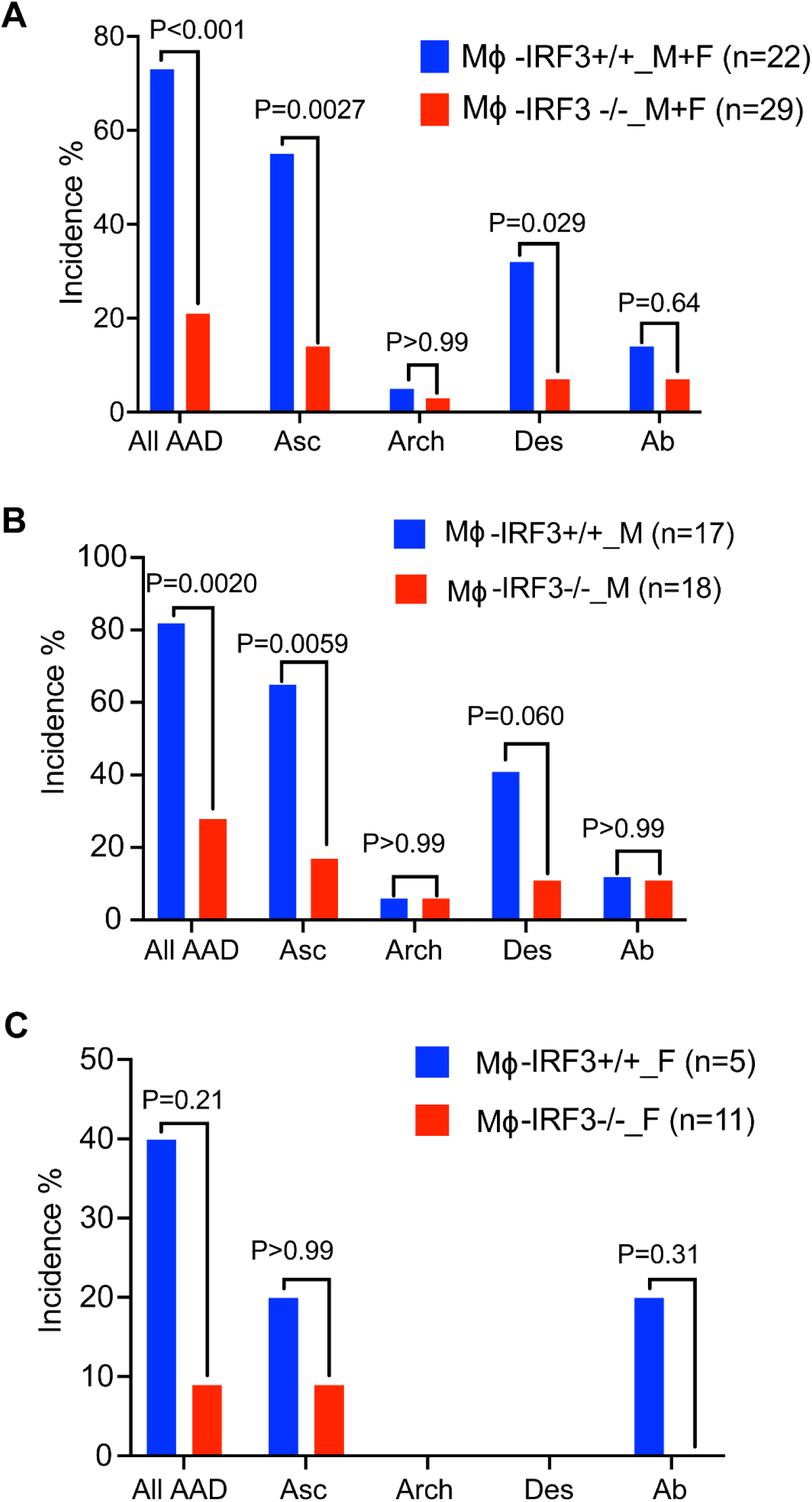
Incidence of aortic dissection in different aortic segments in all MΦ-specific *Irf3* deficient mice **(A)**, and in male **(B)** and female **(C)** mice.

## Nonstandard Abbreviations and Acronyms

Ang II: Angiotensin II
ATAA: ascending thoracic aortic aneurysm
ATAD: ascending thoracic aortic dissection
MΦ: macrophage
IRF3: interferon regulatory factor 3
STING: stimulator of interferon genes
BRG1: brahma-related gene 1
TF: transcription factor
scRNA-seq: single-cell RNA sequencing
scATAC-seq: single-cell sequencing for transposase-accessible chromatin
CUT&RUN-seq: sequencing for cleavage under targets & release using nuclease
IP-MS: immunoprecipitation followed by mass spectrometry

